# Whole-Genome Analysis of Stress Resistance-Related Genes in *Listeria monocytogenes*

**DOI:** 10.1101/2024.01.30.577952

**Authors:** Xin Dou, Yangtai Liu, Efstathios Z. Panagou, Huajian Zhu, Zhuosi Li, Qingli Dong

## Abstract

*Listeria monocytogenes* is a crucial foodborne pathogen with significant public health implications. This study analyzed whole-genome sequences (WGS) of *L. monocytogenes* strains from public databases, examining associations between resistance genes, lineage, strain type, isolation source, and geography. Results revealed that after eliminating duplicates and strains with incomplete WGS, a total of 316 strains were deemed suitable for subsequent analyses. Within these strains, lineages I and II were extensively distributed, predominantly isolated from clinical and food sources. 56.65% of these strains fell into seven major Clonal Complexes (CC), identified by Multilocus Sequence Typing (MLST), correlating significantly with isolation information. Analysis of 46 resistance-related genes showed a high consistency of resistance genes in the same type of strains, hinting at a potential causal chain of ‘habits-foods-environments evolutions’. Moreover, the standard strains exhibit similar gene carriage rates as the sample strains, with multiple variations observed in acid-resistance genes. In conclusion, through a comprehensive analysis of the *L. monocytogenes* genome sequences, this study deepens our understanding of the differences and associations between its lineage, strain typing, isolation sources, geographical distribution, and resistance genes. It has also explored the potential impact of environmental noise on the expression of these genes, offering a scientific foundation for devising more effective prevention and control strategies against *L. monocytogenes*. Future endeavors should further dissect the functions of stress resistance genes and the variations in their expression, with the aim of gaining a deeper insight into the risks posed by *L. monocytogenes* to public health safety.

## 1. Introduction

*Listeria monocytogenes*, a Gram-positive bacterium, can adapt to a variety of environmental conditions (Hu et al. 2023), including a temperature range of 0°C–45°C, pH 4.3–9.6, resistant to high salt concentrations (up to 10.0% NaCl), and low water activity (Aw to 0.90) (Gandhi and Chikindas. 2007a; Wiktorczyk-Kapischke et al. 2021; Wiktorczyk-Kapischke et al. 2023). Its ability to adjust to complex stress responses under varying conditions contributes to its strong adaptability (Martínez-Suárez et al. 2016). *L. monocytogenes* demonstrates an extensive natural distribution, with strains isolated from sources as diverse as water, soil, sewage, decaying vegetation, animal feed, and various species of fish, birds, and mammals (Cavalcanti et al. 2022; Liu et al. 2022; Skowron et al. 2019; Tahir et al. 2022). This wide-ranging presence enhances the likelihood of human exposure and consequent disease manifestation. Food is recognized as the principal vector of human *L. monocytogenes* infections (CDC. 2022), leading to listeriosis which carries a mortality rate of up to 30% (WHO. 2018). It poses a particularly serious threat to the health of the elderly, pregnant women (who can pass it on to their fetuses), newborns, and immunocompromised individuals (Gandhi and Chikindas. 2007a; Pohl et al. 2019).

To further understand the growth characteristics and pathogenic mechanisms of *L. monocytogenes*, it is necessary to conduct a detailed analysis of its strains isolated from different sources using whole-genome sequencing (WGS). In recent years, WGS has played a significant role in the research of *L. monocytogenes*. For instance, Li et al. (2023) performed WGS on 360 strains of *L. monocytogenes* isolated from clinical samples, thoroughly investigating their lineages, serotypes, and Multilocus Sequence Typing (MLST) types. Their research provided valuable information for monitoring listeriosis by thoroughly understanding the genetic diversity among clinically isolated strains. Similarly, Ji et al. (2023) performed WGS on 322 foodborne *L. monocytogenes* strains from 13 regions and four food sources in China, and analyzed their antibiotic resistance and virulence genes in detail. In addition, through WGS, Song et al. (2022) clarified the population structure and characteristics of 207 ST9 *L. monocytogenes* strains from 14 different countries and regions. Meanwhile, Hingston et al. (2017) used WGS to reveal the potential relationship between *L. monocytogenes* genotypes and food-related stress tolerance phenotypes. All these studies demonstrate that WGS can reveal significant differences in genotypes and stress responses among different strains, provide more detailed molecular characteristics, and being helpful to better understand their genetic relationships (Ji et al. 2023). This is of great value to our understanding of the biology and epidemiological research of *L. monocytogenes*.

The survival and proliferation ability of *L. monocytogenes* under various adverse environmental conditions stem from its complex gene regulatory network. These adverse conditions include osmotic pressure, heat, cold, acid, bactericide, and nutritional stress (Hingston et al. 2017). They have prompted *L. monocytogenes* to evolve numerous genomic features to adapt to stress conditions (Wiktorczyk-Kapischke et al. 2021), including stress survival islands (SSI-1 and SSI-2) and other environmental resistance genes, e.g. heat, cold, acid, osmotic, drying, metal, disinfectant, and regulatory genes (Begley et al. 2005; Harter Wagner Eva et al. 2017). Harter (2017) found SSI-1 and SSI-2 in *L. monocytogenes* strains, concluding that the presence of stress survival islands may support the adaptability and persistence of *L. monocytogenes* strains in food processing environments. Omori et al. (2017) analyzed changes in the expression of heat-resistant genes in *L. monocytogenes*, and the results showed that *ClpB* and *ClpE* can enhance the heat resistance of the strains. Liu et al. (2019) analyzed the functional genes regulated by *sigB* and identified 73 stress-related genes regulated by *sigB*, confirming that *sigB* is a very important stress response regulator in *L. monocytogenes*. In summary, *L. monocytogenes*, through corresponding gene regulation, exhibits strong adaptability under diverse adversities, which provides us with the possibility of a deeper understanding and more effective control of listeriosis.

The primary objective of this research is to conduct an in-depth analysis of the WGS of *L. monocytogenes* from public databases, aiming to unveil the disparities and associations between strain resistance characteristics genes and their lineage, strain typing, source of isolation, and geographic region. This includes comparing 316 strains with ATCC standard strains at the genomic level to ascertain the representativeness of these standards in research. The study also investigates the potential influence of environmental noise on the expression of stress resistance genes, providing a crucial reference for the selection of functional genes and research into associated phenotypes. Further comprehension of the genetic diversity among these strains is sought, investigating how these variations impact their resistance, thereby providing valuable insights for the prevention and control strategies of listeriosis.

## 2. Materials and Methods

### 2.1 Information collection

To obtain the whole-genome sequences of the target bacterium *L. monocytogenes*, an initial search was conducted via the official ATCC website (https://genomes.atcc.org/genomes?text=Listeria+monocytogenes) by limiting the collecting date before 15th June 2023. Subsequently, the NCBI website (https://www.ncbi.nlm.nih.gov/) was explored using the keywords: *Listeria monocytogenes* “[Organism] AND “Complete Genome”[Assembly Level] to retrieve all the whole-genome sequences of *L. monocytogenes* available in the database as of the same cut-off date.

To ensure the completeness of genomic sequence and the comprehensiveness of annotations, L. monocytogenes EGD-e was selected as the reference strain. Moreover, the SNPs within EGD-e have been assigned lmo identifiers in major databases, facilitating subsequent research and ensuring the accuracy and reproducibility of experiments. The whole-genome sequence and gene annotation files of the reference strain (*L. monocytogenes* EGD-e) were downloaded from the NCBI website (https://www.ncbi.nlm.nih.gov/genome/159). The collated data were detailed in Supplementary Table 1.

### 2.2 Phylogeny reconstruction based on core genome single nucleotide polymorphisms

The art of Harvest toolkit, *Parsnp*, was employed for core genome alignment of all *L. monocytogenes* target strains and the reference strain EGD-e, to identify single nucleotide polymorphisms (SNPs) within the core genome. To enhance the precision and reliability of the analysis, SNPs clustered within 20 base pairs were eliminated to avoid erroneous SNP calls. The maximum likelihood tree was developed based on the identified core genome SNPs, using a combination of FastTree (version 2.1.11, http://meta.microbesonline.org/fasttree, accessed on 4th May 2020) and the iTOL (version 6.8, https://itol.embl.de/, accessed on 20th April 2023) to form the phylogenetic tree of the strains.

### 2.3 Association analysis between strain typing and isolation information

To estimate the correlation between *L. monocytogenes* strain typing and isolation information (Supplementary Table 2,3), the Chi-Squared test was employed as the primary statistical method. Due to the multiple tests conducted in the study, the Bonferroni correction was applied to adjust the significance threshold, mitigating potential errors (Holm. 1979).

### 2.4 Analysis of stress resistance gene carriage rate

The complete genomic data we obtained was aligned using the Pasteur Institute database (https://bigsdb.pasteur.fr/) to identify the stress resistance genes present across all genomes, and to gather strain typing information along with annotation details (Moura et al. 2016). To ensure precise identification of stress resistance genes, we set the parameters to include sequences with >95% nucleotide identity and >90% sequence coverage (Nelson et al. 2004). Following this, R software (version 4.2.1, https://cran.r-project.org/, accessed on 18th June 2022) was employed to visually analyze the detection status of stress resistance genes. An unweighted pair group method with arithmetic mean (UPGMA) was applied to generate a heat map for hierarchical clustering.

### 2.5 Mutation analysis of stress resistance genes

The sequence of the *L. monocytogenes* standard strain EGD-e was downloaded from the NCBI database (https://www.ncbi.nlm.nih.gov/) as a reference sequence. Predicted stress resistance gene sequences were translated into protein sequences using the Dnastar 7.1 software (DNASTAR Inc., Madison, WI, USA) and aligned with the reference protein sequence to predict whether any mutations occurred in the stress resistance genes.

## 3. Results

### 3.1 Data acquisition and strain selection

A total of 340 strains with uploaded whole-genome sequences were obtained from the search of whole-genome sequences of *L. monocytogenes* in the NCBI and ATCC databases. Within these, the ATCC database contained 29 standard strains of *L. monocytogenes*, while the NCBI had 311 strains with complete genome sequences. To ensure data quality and accuracy, strains were refined by removing duplicates and those with incomplete whole-genome sequences, following the flow shown in Fig. 1 and resulting in 316 analyzable *L. monocytogenes* strains.

**Fig. 1.**
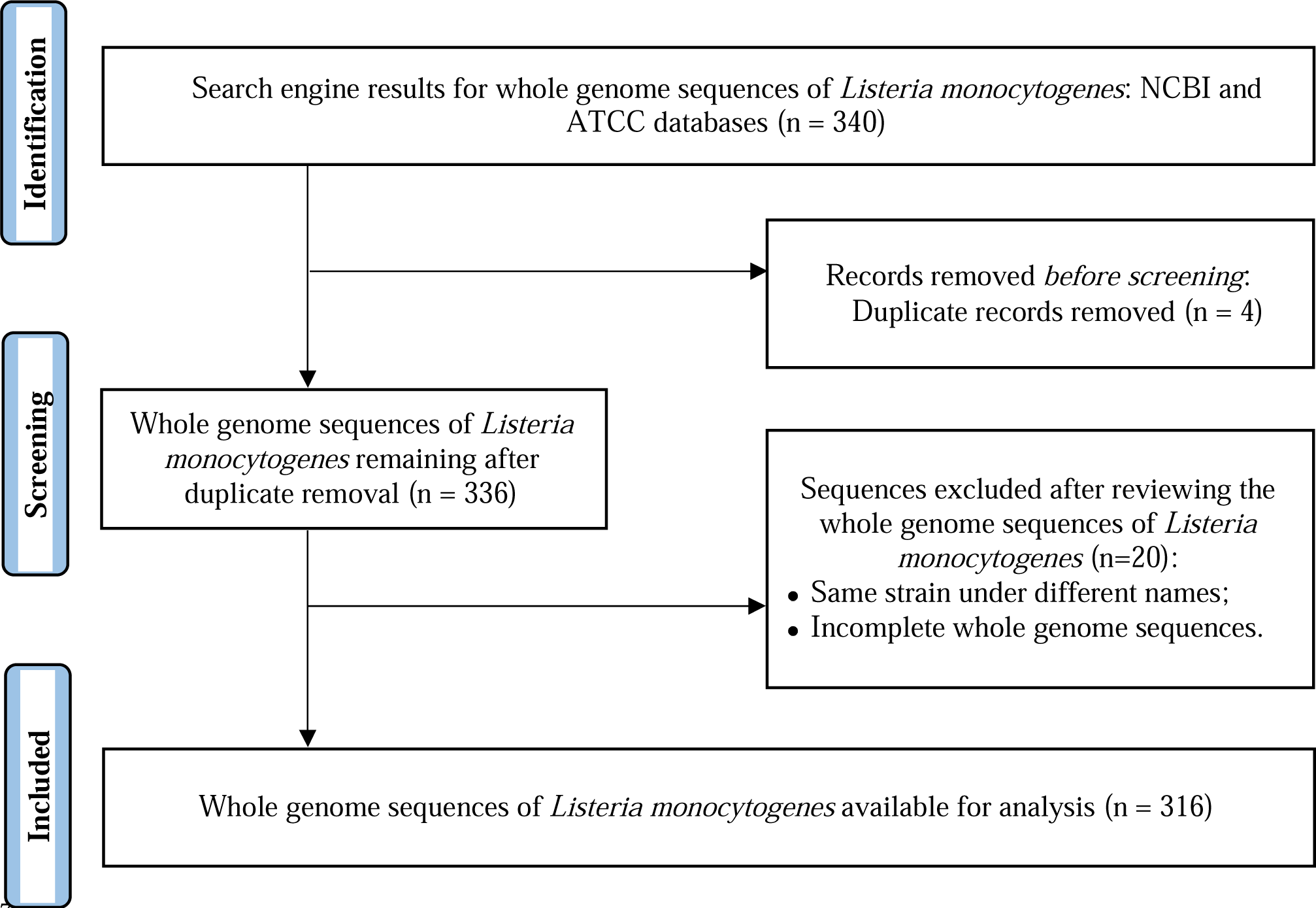
Flowchart of obtaining whole genome sequence of *Listeria monocytogenes*

### 3.2 Distribution of *L. monocytogenes*

Isolation source and regional data for *L. monocytogenes* were obtained from the database. As depicted in Fig. 2(A), the most abundant category of the collected *L. monocytogenes* strains was isolated from clinical cases, comprising 40.1% of the total. The second most common source was food, accounting for 30.5% of the isolates. This distribution underscored food as a predominant route of transmission for *L. monocytogenes*. Once ingested via contaminated food, the bacteria could develop into clinical disease, thereby explaining its frequent isolation from clinical samples.

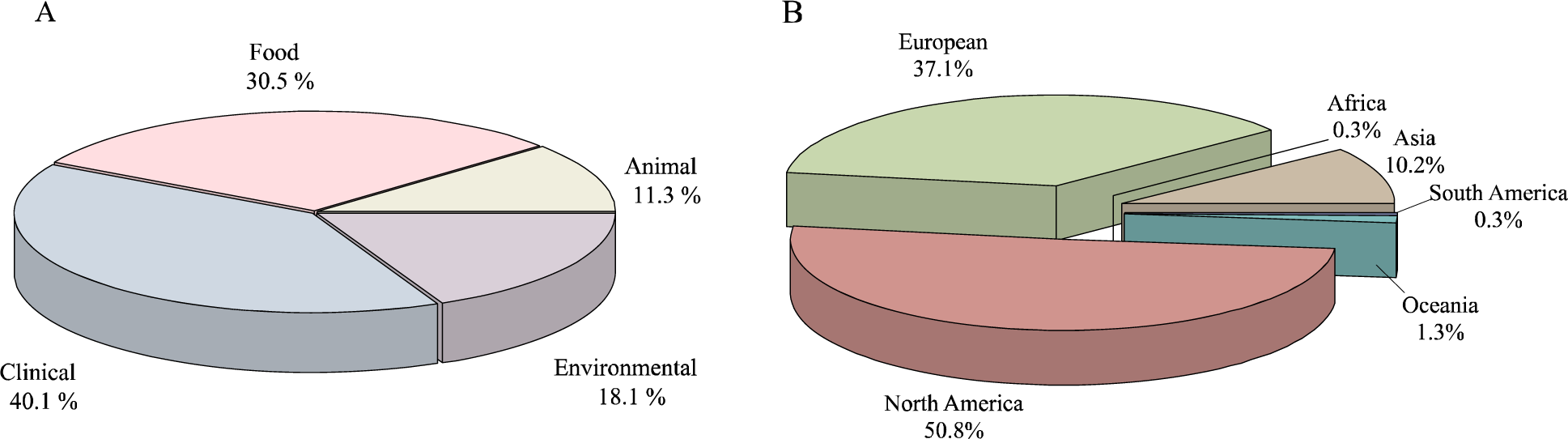

In the geographical distribution diagram, shown in Fig. 2(B), the prevalence of *L. monocytogenes* was more pronounced in Europe and North America, while in regions such as Asia, Africa, Oceania, and South America, the distribution was comparatively sparse. This disparity in geographical distribution could likely have been attributed to two principal factors: Dietary habits varying across regions might have influenced the transmission patterns of *L. monocytogenes*, thereby altering its distribution across different geographical areas; and the usage of the database and the geographic distribution of data contributors could have introduced a bias, potentially affecting the observed distribution of the bacteria.

### 3.3 Characteristics of *L. monocytogenes*

*L. monocytogenes* was categorized based on the lineage, isolation source, and isolation region, and underwent MLST typing to construct the phylogenetic tree shown in Fig. 3. The lineage classification of *L. monocytogenes* serotypes, collected from the database, mainly included four types: Lineage I, II, III, and IV. Notably, Lineages I and II were most prevalent in the sample pool, indicating a potentially rich presence of these strains in nature. Furthermore, Lineage I strains were predominantly located at the root of the phylogenetic tree, signifying their ancient origin. The phylogenetic tree also suggested that Lineage I strains were the ancestors of the other lineages (II, III, and IV), hence they might have evolved into new lineages through a series of evolutionary mechanisms, such as gene mutations, recombination, and horizontal gene transfer. It was noteworthy that the ATCC standard strains were predominantly found within lineages I and II, which further underscores the abundance and widespread distribution of strains from these lineages in the natural environment.

**Fig 3.**
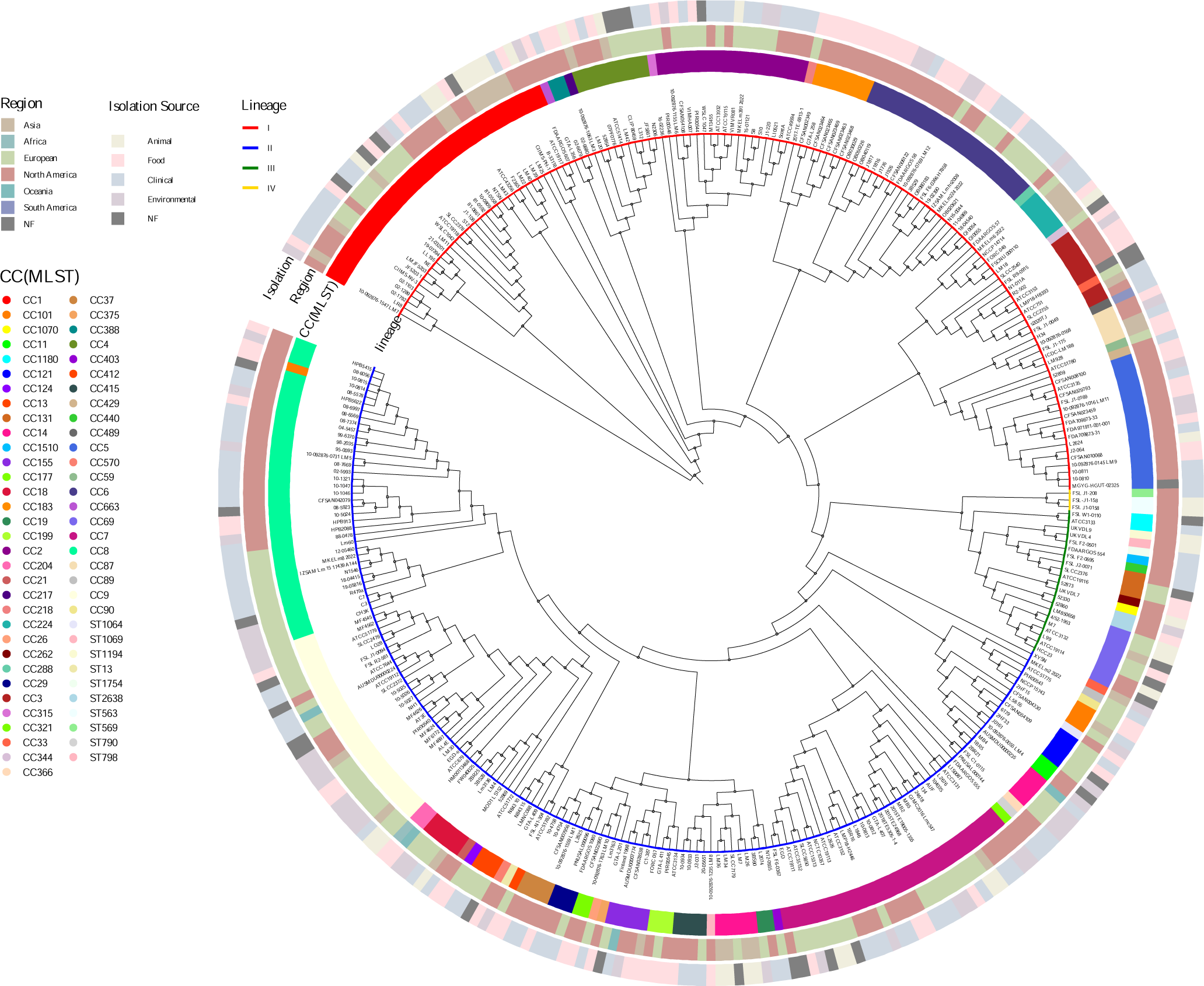
Phylogenetic tree of L. monocytogenes illustrating lin eages, isolation regions, sources, and strain types.

### 3.4 Correlation analysis between strain typing and isolation information

A deeper analysis was conducted on the association between the strain types and isolation data of *L. monocytogenes*. Of the aforementioned 316 *L. monocytogenes* strains, they were classified into 67 distinct CC types. Stacked diagrams were crafted based on the MLST (CC type) classification of *L. monocytogenes* with varying isolation information (Fig 4a, 5a), among which the seven most prevalent CC types accounted for 56.65% of all strains in this study. Additionally, the frequency distribution of these distinct CC types varied slightly, with respective proportions of: CC8 (11.08%), CC1 (10.76%), CC7 (9.18%), CC9 (7.91%), CC6 (6.96%), CC2 (5.70%), and CC5 (5.06%). Subsequent analyses examined the correlation between strain types and distinct isolation sources, as well as strain types and varying isolation regions, revealed predominant CC types in different isolation sources (Fig 4b; Supplementary Table 2). CC183 was significantly associated with food sources (*p* < 0.01); CC8 showed a certain correlation with clinical sources (*p* < 0.05); CC9 was significantly associated with environmental sources (*p* < 0.01); and CC14 with animal sources (*p* < 0.01). Further analysis of the relationship between strains and their geographic origins also unveiled specific patterns (Fig 5b; Supplementary Table 2): CC9 exhibited a notable correlation with Europe (*p* < 0.05), while CC1 and CC5 were significantly associated with North America (*p* < 0.05 and *p* < 0.01, respectively). Furthermore, the most significant associations with Asia were observed in CC14 (*p* < 0.01) and CC224 (*p* < 0.001). In summary, this data vividly portrayed the distribution patterns of L*. monocytogenes* in various isolation contexts, underscoring the significance of specific CC types in certain sources and regions.

**Fig. 4.**
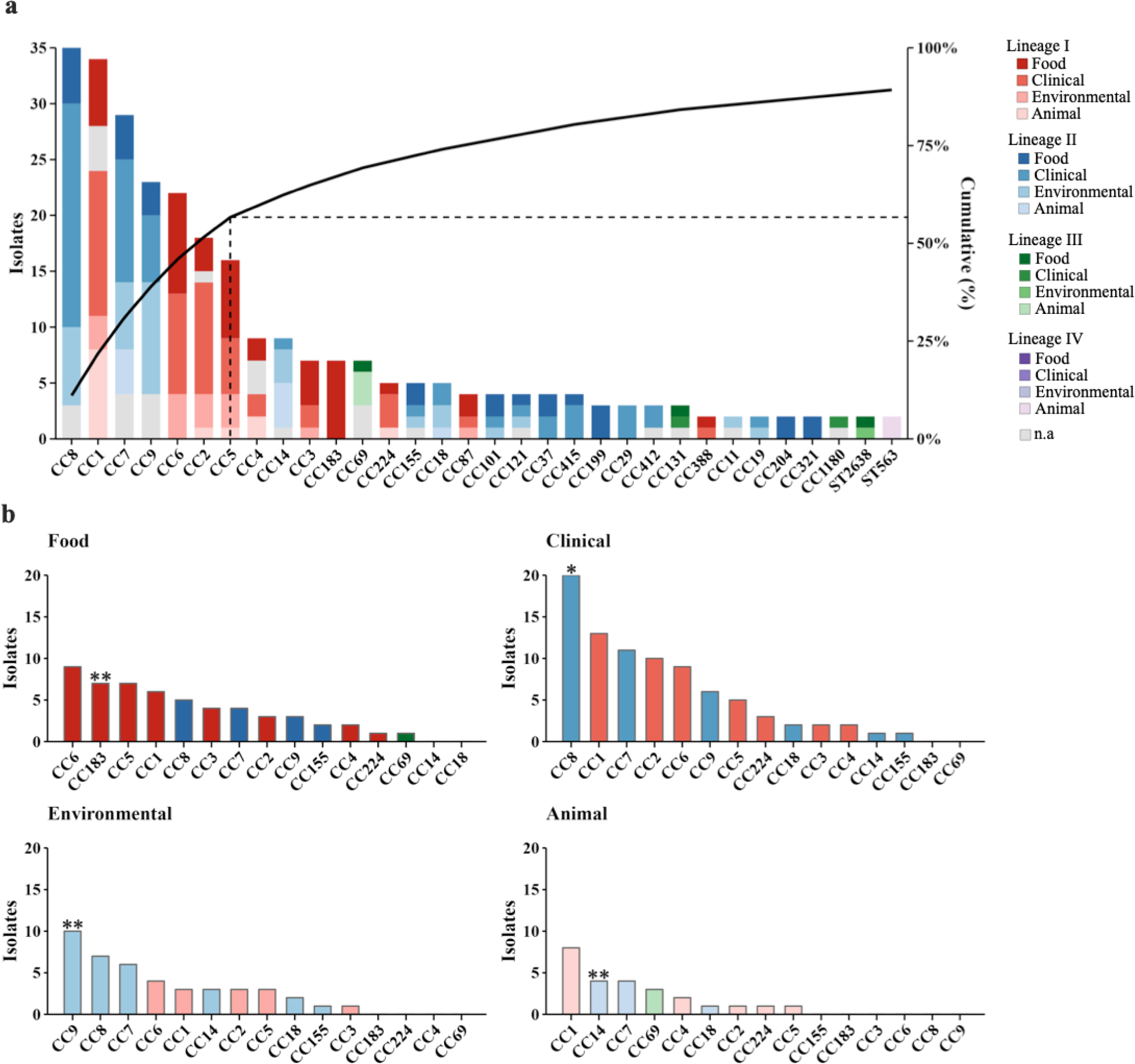
Prevalence and distribution of MLST (CC type) in different isolation sources

**Fig. 5.**
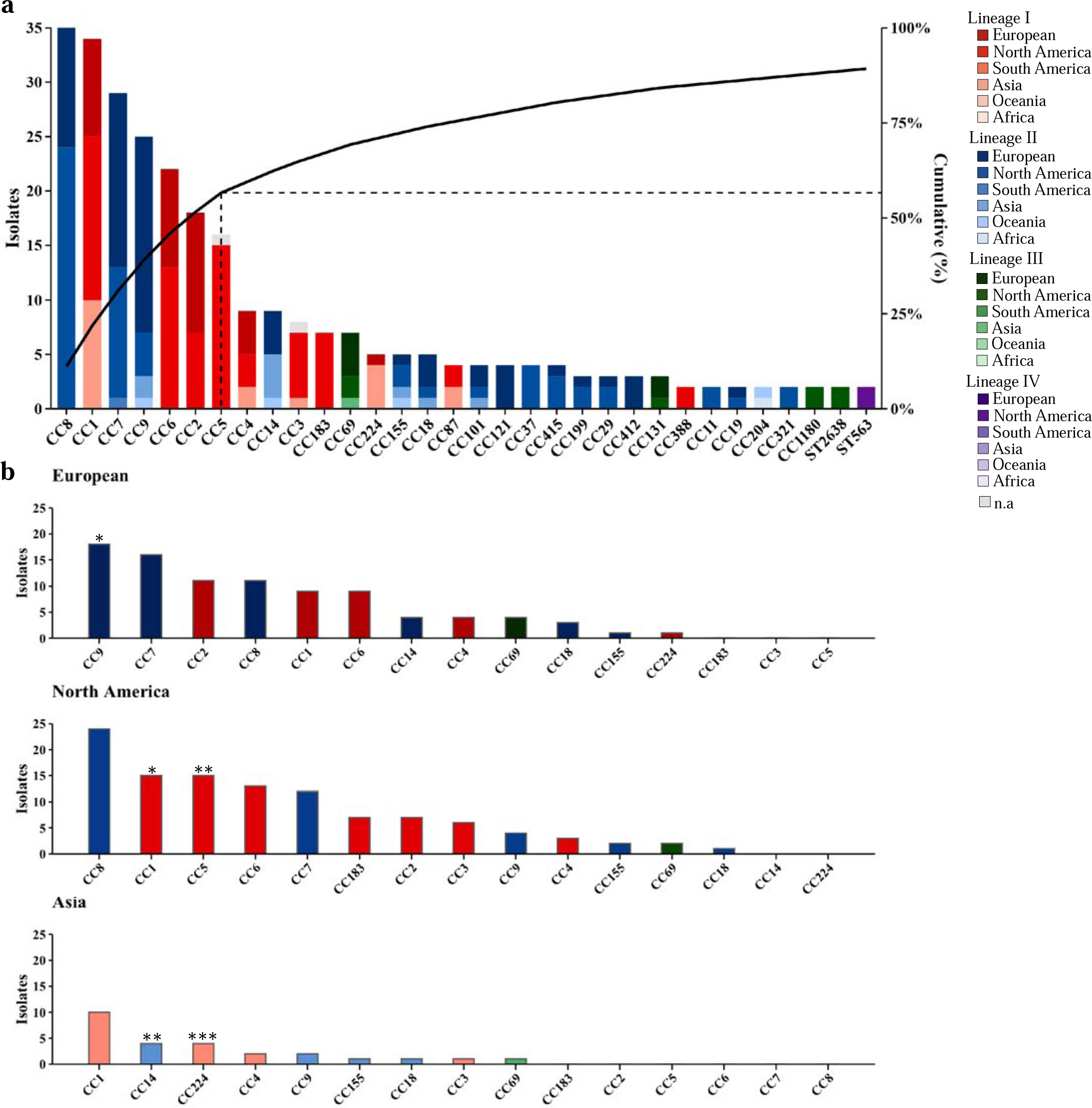
Prevalence and distribution of MLST (CC type) in different isolation region

### 3.5 Stress resistance gene carriage rate

According to literature and gene annotation information, a total of 46 function genes related to the resistance traits of *L. monocytogenes* were extracted for subsequent analysis, as summarized in Table 1. These genes played a critical role in the survival and adaptation of *L. monocytogenes* in various environments, enabling them to resist diverse adverse conditions such as high temperatures, cold, acidic environments, hyperosmotic environments, desiccation, metals, and biocides (Holt et al. 2015).

**Table 1.** Compilation of Resistance Genes Associated with *L. monocytogenes* Across Various Stress Factors.

| Category | Related genes | Reference |
| --- | --- | --- |
| SSI-1 | *lmo0444, lmo0445, pva, lmo0447, lmo0448* | (Hilliard et al., 2018) (Ryan et al., 2010) |
| SSI-2 | *lin0464, lin0465* | (Harter, Wagner, et al., 2017) |
| Heat Resistance | *dnaK, clpB, clpE, clpP, mcsB, ctsR* | (Fang et al., 2016; Omori et al., 2017) |
| Cold Resistance | *cspB, cspD, gbuA, gbuB, gbuC, betL, opuCA, opuCB, opuCC, opuCD, lisK, lmo1078* | (Smith et al., 2019; Wemekamp-Kamphuis et al., 2004) |
| Acid Resistance | *arcA, arcB, arcC, arcD* | (Chen et al., 2010; Hingston et al., 2017; Liu et al., 2019) |
| Osmotic Resistance | *gbuA, gbuB, gbuC, betL, opuCA, opuCB, opuCC, opuCD* | (Kang et al., 2019) |
| Desiccation Resistance | *fliP, flhB, flgD, flgL, motA, motB, fliM, fliY* | (Casey et al., 2014) |
| Resistance to Metals and Disinfectants | *bcrA, bcrB, bcrC, emrE, emrC, emrA, cadA* | (Cherifi et al., 2018; Kovacevic et al., 2016) |
| Regulation | *sigB, prfA, prfB* | (Ait-Ouazzou et al., 2012; Harter, Wagner, et al., 2017) |

Simultaneously, based on the gene sequence of the standard strain EGD-e, an exhaustive sequence comparison of these 46 resistance trait genes was conducted among the 316 sample strains and the ATCC standard strains. Utilizing the Pasteur Institute database (https://bigsdb.pasteur.fr/), we thoroughly analyzed the carriage rate of the resistance trait genes in these strains and explored the gene differences between the 316 sample strains and the ATCC standard strains. As shown in Fig 6, the majority of the stress response genes’ carriage rates in the 316 sample strains paralleled those in the reference strain. Regarding individual stress response genes, it was observed that most could find a counterpart in both the reference strain and the 316 sample strains. The similar carriage rates of these genes in both the reference and the sample strains suggested that the ATCC standard strains were typical representatives.

**Fig. 6.**
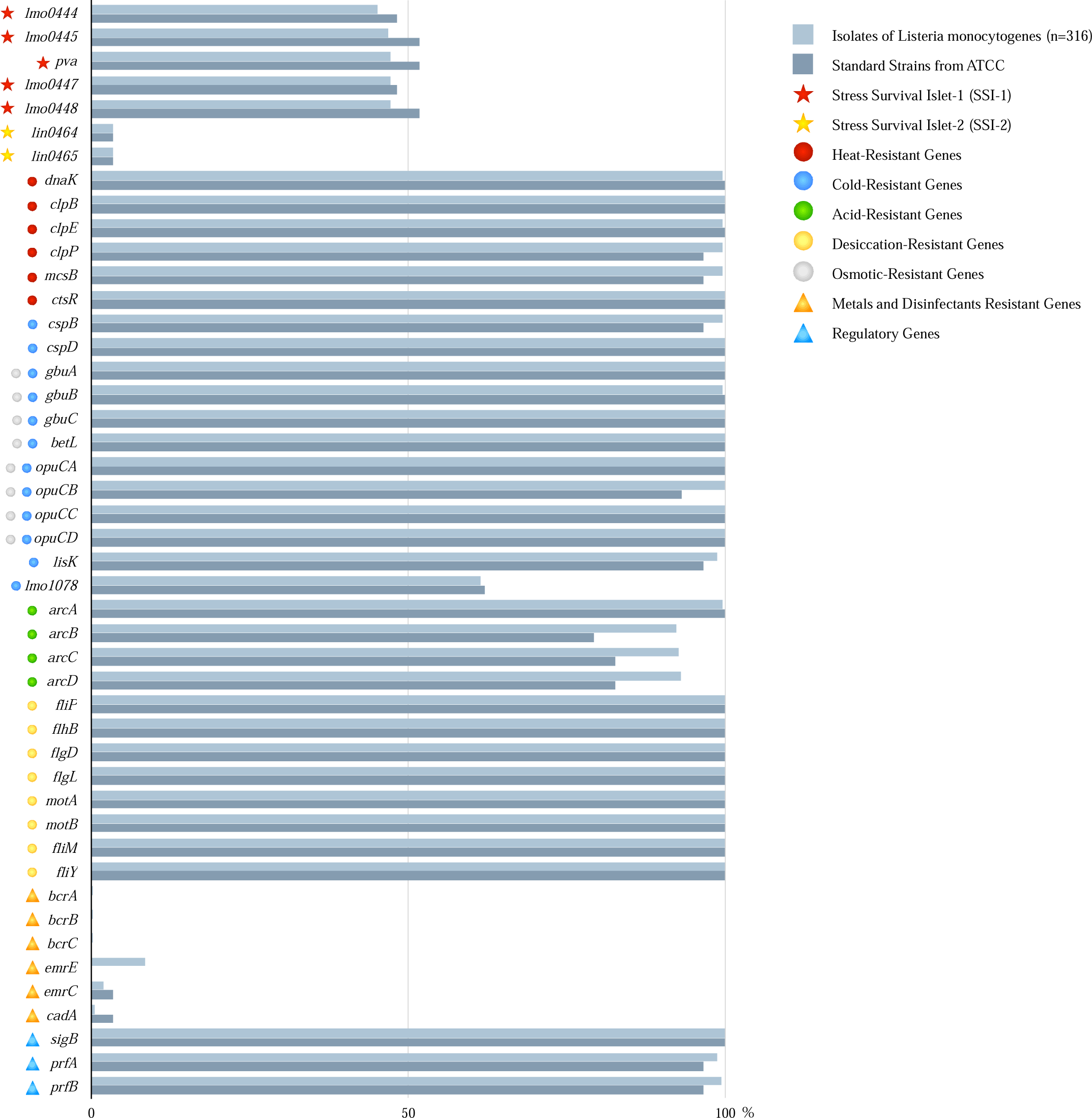
Carrying rate of *L. monocytogenes* resistance characteristic gene

However, the raw data comparison further revealed some subtle differences. While the ATCC standard strains essentially encompassed all resistance trait genes from the 316 sample strains, three genes could not be found in the ATCC standard strains. Notably, only one strain among the 316 sample strains carried these three genes. This indicated that while the ATCC standard strains incorporated most of the resistance trait genes, some were still not included.

Furthermore, a deeper dive into the raw data to contrast the carriage of resistance genes in various *L. monocytogenes* strain types uncovered that acid resistance genes showed the most pronounced variation, with other resistance genes displaying minor differences. Concurrently, strains of the same CC type demonstrated a high degree of consistency in their repertoire of stress response genes. Within a single CC type, strains isolated from the same source exhibited nearly identical profiles of these genes. Across different isolation sources, strains derived from animals generally harbored genes for heat, cold, desiccation resistance, and regulatory genes. Similarly, strains from clinical, environmental, and food sources typically possessed genes for heat, cold, acid, desiccation resistance, and regulatory genes. Such a gene profile facilitated the survival of *L. monocytogenes* under challenging conditions, explaining their detection across varied isolation sources, a trend also mirrored in the reference strain. In general, the ATCC standard strains demonstrated typical in terms of carriage rate and gene diversity. Although some genes were not encompassed, on a broader scale, the ATCC standard strains still represented the distribution and existence of the majority of the resistance trait genes.

### 3.6 mutation analysis of stress resistance genes

Due to the typical ATCC standard strains, they were selected for detailed investigation, specifically examining the carriage and variations of stress-resistance genes within *L. monocytogenes* ATCC strains. As Fig. 7 illustrates, genes resistant to metals and germicides were not detected in the majority of ATCC standard strains. In the context of stress islands, the SSI-1 stress island was detectable in half of the ATCC standard strains, whereas the SSI-2 stress island was detected only in ATCC51782. Concurrently, heat resistance, cold resistance, acid resistance, high osmotic resistance, desiccation resistance, and regulatory genes were ubiquitously present across all ATCC strains. Notably, desiccation resistance genes could be entirely detected in all ATCC standard strains. Furthermore, eight genes (*gbuABC*, *betL*, and *opuCABCD*) demonstrated both high osmotic and cold resistance and were detected in all ATCC standard strains. However, among all these genes, acid resistance genes exhibited the greatest variability in detection rates across different ATCC standard strains.

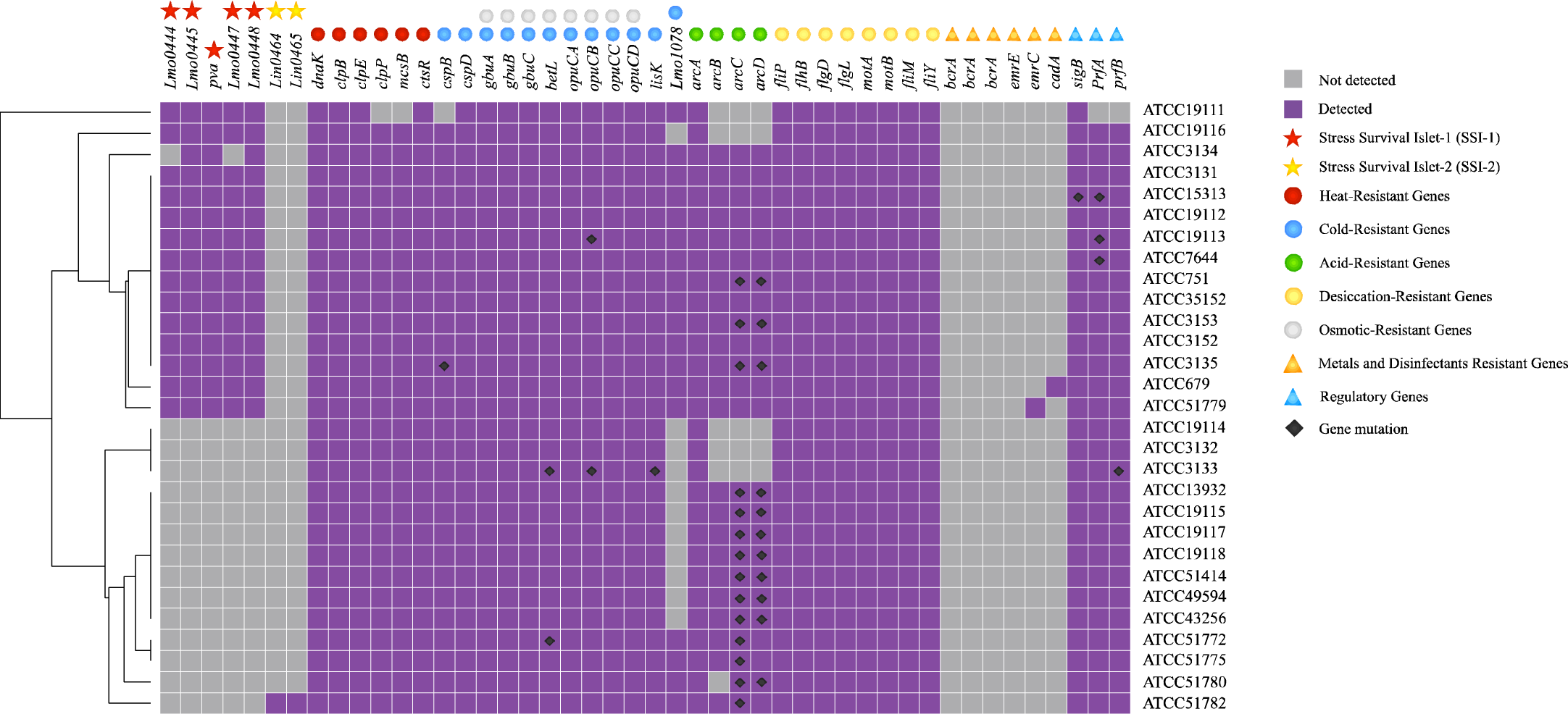

To further elucidate the variations in these resistance genes, mutations within the gene sequences were analyzed. Our results revealed that out of these 46 genes, 18 ATCC standard strains had experienced gene sequence mutations. Among these strains, ATCC3133 exhibited the highest number of mutated genes, with four genes (*betL*, *opucB*, *lisK*, and *prfB*) undergoing mutations. At the gene level, the acid resistance genes *arcC* and *arcCD* were the most commonly mutated genes in ATCC standard strains, with mutation rates of 60.87% and 47.83%, respectively. In summary, differences existed in the carriage and mutations of stress-resistance genes among ATCC standard strains. These differences held significant implications for understanding the stress resistance mechanisms of *L. monocytogenes*.

## 4. Discussion

*L. monocytogenes* is a significant foodborne pathogen, causing listeriosis, a substantial public health problem globally (Cossart. 2011). While many studies have delved into the genomic sequences of *L. monocytogenes*, the relationships between the strain’s lineage, typing, isolation source, and geographic region of isolation remained ambiguous. In this research, a thorough analysis of the genomes of 316 *L. monocytogenes* strains was conducted, shedding light on the correlations and variations between the strain’s stress response genes, lineage, strain typing, isolation source, and geographic area of isolation. Moreover, it examines the potential effects of environmental noise on the expression of stress resistance genes, thereby aiding in the comprehension of the adaptability and transmission routes of *L. monocytogenes*.

Extensive phylogenetic and subtyping research revealed that *L. monocytogenes* exhibited a distinct population structure, encompassing at least four different evolutionary lineages: I, II, III, and IV (Liu et al. 2006). Within these lineages, Lineages I and II boasted the most typical strains and had been associated with the majority of human clinical cases (den Bakker et al. 2008; Orsi et al. 2011). In our study, we classified 316 strains of *L. monocytogenes* into these lineages and found a broad representation of lineages I and II in all the samples. Gray et al. (2004) analyzed 502 *L. monocytogenes* isolates from food and 492 from clinical, respectively, and found that Lineage I and III were mainly derived from clinical samples, while

Lineage II was mainly derived from food samples. However, they detected only a small proportion of lineage III strains in both food and clinical samples, thus hypothesizing that lineage III strains struggled to survive in food processing or storage environments. Notably, our research also demonstrated a lower prevalence of lineage III strains in our samples, which aligned with the findings of Gray et al. (2004). As for lineage IV, Orsi et al. (2011) argued that while it was most frequently detected in animal isolates and can also be isolated from human cases, its limited number of isolates means that it is often not extensively discussed in some studies.

In addition to phylogenetic distribution, we further developed into the variations of *L. monocytogenes* in terms of isolation sources and geographic locations. The *L. monocytogenes* isolates obtained from the database in our study were divided into four major categories, mainly including clinical, food, animal, and environmental sources, among which isolates from clinical and food sources accounted for the largest proportion. As *L. monocytogenes* is a foodborne pathogen, diseases are usually triggered after human consumption of food, which has been validated in multiple studies (Maury et al. 2016). Chen et al. (2016) summarized outbreaks of listeriosis on a global scale, discovering that outbreaks were mostly food-related, and corresponding *L. monocytogenes* was detected in the clinic, further validating our findings. Moreover, a large number of *L. monocytogenes* were also isolated from animal and environmental sources, which are equally significant causes of disease outbreaks.

In the analysis of geographical distribution differences, it was reasonably posited that dietary habits might influence the transmission patterns of *L. monocytogenes* consequently altering its distribution across various regions. Some studies have shown that dietary habits indeed could influence the incidence of listeriosis. For instance, consumption of undercooked meats, preparing raw foods, or making soft cheese at home have been identified as risk factors for listeriosis (Silk et al. 2014). Additionally, regional outbreaks of listeriosis, such as the Boston cheese incident (Pightling et al. 2014) and packaged salad outbreaks in the United States (Centers for Disease Control and Prevention. 2016), suggest that regional dietary preferences could contribute to the occurrence of listeriosis. However, the acquisition and analysis of research data are often influenced by geographical and temporal factors, and a substantial portion of the data, due to its confidential nature, is not uploaded to public databases. Therefore, these findings necessitate further validation. Despite these limitations, a deep understanding of the geographic distribution of *L. monocytogenes* and dietary exposure can undoubtedly aid in better prevention and control of listeriosis.

Further analysis of the association between strain typing and isolation information revealed that 56.65% of the 316 *L. monocytogenes* strains in this study belong to seven predominant CC types. The pronounced clustering suggests these primary CC types might possess greater adaptability or transmission capability in certain environments or conditions. Specifically, CC8, accounting for 11.08%, is closely linked to clinical sources, a finding corroborated by Maury et al. (2016) who also observed a significant correlation between CC8 and clinical origins, especially bacteremia. CC183’s strong association with food sources indicates this CC type may possess specialized adaptive mechanisms during food processing, storage, or transmission. The potent link between CC14 and animal origins implies it has unique survival capabilities within animal hosts. Geographically, the associations of CC9 with Europe, CC1 and CC5 with North America, and CC14 and CC224 with Asia, suggest strain distribution is influenced by local culture, economic activities, food consumption habits, and public health initiatives, underscoring the significance of regional factors. However, data acquisition and analysis are often influenced by geographic and temporal factors. A substantial portion of the data, considered confidential, isn’t uploaded to public databases, necessitating further validation of these findings. Despite these limitations, analyzing the origins and regions of *L. monocytogenes* can assist in better controlling the outbreak of listeriosis.

Distinct dietary habits shape the selection, processing, and storage of food, leading to variations in its physical and chemical properties, e.g., pH, salinity, and water activity. These changes may apply evolutionary pressure to *L. monocytogenes*, driving the emergence of corresponding resilience (Gandhi and Chikindas. 2007b). Stress resistance genes play pivotal roles in *L. monocytogenes*, endowing the bacteria with abilities to thrive and reproduce under various environmental pressures (Chaturongakul and Boor. 2004). As evident from Figure 4, different *L. monocytogenes* types display variations in the carriage of resistance genes, with acid resistance genes showing the most significant differences. Furthermore, strains of the same CC type consistently harbor similar stress resistance genes, reflecting the gene’s stability and the evolutionary pressures unique to that CC type. This consistency aids predictions in clinical or public health contexts, as knowledge of a strain’s CC type can give insights into its resilience attributes. Concurrently, analyzing stress resistance genes in *L. monocytogenes* across different isolation sources provides insights into their adaptability. For instance, strains from animal origins tend to carry genes for heat and cold resistance, likely correlated with animal body temperatures and external environmental fluctuations.

Further examination of gene differences among the collected strains revealed that the carriage rate for genes in the Stress Survival Island (SSI)-1 is approximately 50%, with no observed gene mutations. Typically, SSI-1 enhances the tolerance of *L. monocytogenes* in food processing environments and plays a significant role under stress conditions such as low pH and high salinity (Ryan et al. 2010). Moreover, SSI-1 is closely associated with the biofilm formation of *L. monocytogenes* under environmental stress conditions (Keeney et al. 2018). Ryan et al. (2010) analyzed the stress resistance characteristics of SSI-1 genes in the EGD-e strain, indicating that SSI-1 might enhance the adaptability to food environments. However, in Hingston’s phenotype validation of other strains (Hingston et al. 2017), no significant stress tolerance differences were observed in strains possessing SSI-1. Hence, further in-depth studies are warranted, employing methods such as gene knockout.

Other genes, including heat, cold, acid, osmotic, desiccation, and regulatory genes, are commonly present in the 316 strains of *L. monocytogenes* and the ATCC standard strain. In heat resistance gene-related research, Omori et al. (2017) powerfully validated the participation of heat resistance genes in *L. monocytogenes* expression. In cold resistance genes, the *lisK* gene encodes arginine kinase, enabling *L. monocytogenes* to grow at low temperatures (Pöntinen et al. 2015). The functions of acid resistance (Hingston et al. 2017)), osmotic resistance (Kang et al. 2019), desiccation resistance (Hingston et al. 2017), and regulatory genes (Chaturongakul and Boor. 2004) were also confirmed in the study, further illustrating their significant impact on enhancing strain tolerance. However, in the gene variation results of the ATCC standard strain, variations in acid-resistance genes were common. Therefore, this part still requires more in-depth genetic information analysis (such as SNPs) to better understand the influence of gene variation on the growth and propagation of *L. monocytogenes*.

From the discussions above, it’s evident that the variance of resistance genes in *L. monocytogenes* across different strain types and isolation data can illuminate the bacterium’s evolutionary adaptation capabilities under varying conditions. Existing research has already demonstrated that *L. monocytogenes* in foods typically stored in high-salt or low-temperature environments may possess heightened salt or cold resistance compared to those in other conditions (Elena and Lenski. 2003; Nightingale et al. 2004). The strain type is somewhat linked with both the source of isolation and the geographical area, and resistance genes within a strain type exhibit consistency. This suggests that prolonged environmental adaptation can lead to specific genotypic evolution in *L. monocytogenes* (Durack et al. 2013), hinting at a potential causative chain: “consumption habits-food-food environment-evolution of *L. monocytogenes*.” These genes are also prevalent in standard strains, with variations in acid resistance genes aligning with sample strains, making standard strains highly typical for studying stress resistance attributes. However, it’s noteworthy that of the 53,186 *L. monocytogenes* strains uploaded to the NCBI database, only 0.58% possess complete genomic sequences, indicating ample scope for further research.

While significant variability and divergence have been noted in the resistance genes across different strains, identical genes within strains exhibit notable effects on their resilience due to random fluctuations in gene expression—termed ‘noise’. Studies have suggested that intrinsic noise, such as stochastic events during transcription and translation, and extrinsic noise, induced by environmental changes affecting gene expression, can variably impact a strain’s resistance traits (Raser and O’Shea. 2005). This is particularly pronounced in foodborne pathogens, where extrinsic noise may play a larger role in determining cellular phenotypes (Rosenfeld et al., 2005). Research by Suzuki et al. (2014) using expression profiles to predict antibiotic resistance has indicated that even minor variations in gene expression levels can lead to significant changes in resistance, underscoring the impact of gene expression noise on microbial resistance. Erickson et al. (2017) analyzed the variability of gene expression in bacteria undergoing adaptive evolution, finding that expression variability correlates with a population’s tolerance to stress, reflecting the bacteria’s capacity to adapt to pressures, including the influence of gene expression noise. Thus, a deeper investigation into the relationship between gene expression noise and strain resistance is not only crucial for understanding how genes maintain functionality in diverse environments but also for developing strategies to prevent and control the transmission of foodborne diseases.

## 5. Conclusion

In this study, we screened the full genome sequences of *L. monocytogenes* from public databases, obtained 316 strains with complete full genome sequences, and further performed comparative analyses. The results reveal that Lineage I and II strains are most widely distributed in the natural world, with clinical cases and food serving as their primary isolation sources. The geographical distribution of strains exhibits notable regional disparities. Based on the correlation results between clonal complex (CC) typing and isolation information, different CCs are closely related to their respective sources. In the evaluation of 46 stress-resistance genes, strains of the same type demonstrated high consistency in resistance gene carriage, suggesting a potential causative chain encompassing ‘habits-foods-environments evolutions’. Furthermore, the gene carriage rate in the standard strains closely mirrored that of the sample strains, underscoring the typical representativeness of the ATCC standard strains. Further analysis of the stress-resistance genes in ATCC standard strains revealed a prevalent variation in acid-resistance genes. The impact of this genetic variation on gene functionality warrants further exploration. Beyond genetic variations, the potential impact of noise in gene expression also warrants further attention. In summary, this research provides a comprehensive analysis of the *L. monocytogenes* genome sequences, contrasting strains based on lineage, CC, isolation source, geographical distribution, and stress-resistance genes. This contributes to understanding the resistance mechanisms of *L. monocytogenes* and offers insights for risk management strategies.

## Supporting information

https://docs.google.com/spreadsheets/d/1aSnTY5T0N0mPPYv5xO2CIRNgZnopPq4r/edit?usp=drive_link&ouid=104369327746861774634&rtpof=true&sd=true

https://docs.google.com/spreadsheets/d/1J1F5xvIJ3JBPXfz4FaxnzVAqYkzTecyl/edit?usp=drive_link&ouid=104369327746861774634&rtpof=true&sd=true

https://docs.google.com/spreadsheets/d/1zNzC3k_EQRzeQYRjVc9TfpaE3JvRY5mS/edit?usp=drive_link&ouid=104369327746861774634&rtpof=true&sd=true

## Data availability

Data will be made available on request.

## Acknowledgements

This research was funded by the National Natural Science Foundation of China (32102095), Jiangsu Provincial Science and Technology Plan Special Fund (BZ2023004), and NMPA Key Laboratory for Testing Technology of Pharmaceutical Microbiology (2023-WSW-02).

## Reference

Ait-Ouazzou A, Manas P, Condón S, Pagán R, García-Gonzalo D (2012) Role of general stress-response alternative sigma factors σ^S^ (RpoS) and ^B^ (SigB) in bacterial heat resistance as a function of treatment medium pH. Int J Food Microbiol. 153(3):358–364. 10.1016/j.ijfoodmicro.2011.11.027

Begley M, Sleator Roy D, Gahan Cormac GM, Hill C (2005) Contribution of three bile-associated loci, *bsh*, *pva*, and *btlB*, to gastrointestinal persistence and bile tolerance of *Listeria monocytogenes*. Infect Immun 73(2): 894–904. 10.1128/iai.73.2.894-904.2005

Casey A, Fox EM, Schmitz-Esser S, Coffey A, McAuliffe O, Jordan K (2014) Transcriptome analysis of *Listeria monocytogenes* exposed to biocide stress reveals a multi-system response involving cell wall synthesis, sugar uptake, and motility. Front Microbiol 5(2): 68. 10.3389/FMICB.2014.00068

Cavalcanti AAC, Limeira CH, Siqueira INd, Lima ACd, Medeiros FJPd, Souza JGd, Medeiros NGdA, Oliveira Filho AAd, Melo MAd (2022) The prevalence of *Listeria monocytogenes* in meat products in Brazil: A systematic literature review and meta-analysis. Res Vet Sci 145(7):169–176. 10.1016/j.rvsc.2022.02.015

Centers for Disease Control and Prevention [CDC] (2022) Listeria Outbreaks. Listeria Outbreaks Homepage | CDC. (Accessed 01 March 2023).

Chaturongakul S, Boor KJ (2004) RsbT and RsbV contribute to σ^B^-dependent survival under environmental, energy, and intracellular stress conditions in *Listeria monocytogenes*. Appl Environ Microbiol 70(9):5349–5356. 10.1128/aem.70.9.5349-5356.2004

Chen J, Chen Q, Jiang J, Hu H, Ye J, Fang W (2010) Serovar 4b complex predominates among *Listeria monocytogenes* isolates from imported aquatic products in China. Foodborne Pathog Dis 7(1):31–41. 10.1089/fpd.2009.0353

Chen Y, Gonzalez-Escalona N, Hammack Thomas S, Allard Marc W, Strain Errol A, Brown Eric W (2016) Core genome multilocus sequence typing for identification of globally distributed clonal groups and differentiation of outbreak strains of *Listeria monocytogenes*. Appl Environ Microbiol 82(20):6258–6272. 10.1128/AEM.01532-16

Cherifi T, Carrillo C, Lambert D, Miniaï I, Quessy S, Larivière-Gauthier G, Blais B, Fravalo P (2018) Genomic characterization of *Listeria monocytogenes* isolates reveals that their persistence in a pig slaughterhouse is linked to the presence of benzalkonium chloride resistance genes. BMC Microbiol 18(1):1–13. 10.1186/s12866-018-1363-9

Control CfD, Prevention (2016) Multistate outbreak of listeriosis linked to packaged salads produced at Springfield. In Ohio Dole Processing Facility (Final Update). (Accessed 01 March 2023).

Cossart P (2011) Illuminating the landscape of host–pathogen interactions with the bacterium *Listeria monocytogenes*. Proc Natl Acad Sci USA 108(49):19484–19491. 10.1073/pnas.1112371108

den Bakker HC, Didelot X, Fortes ED, Nightingale KK, Wiedmann M (2008) Lineage specific recombination rates and microevolution in *Listeria monocytogenes*. BMC Evol Biol 8(1):277. 10.1186/1471-2148-8-277

Durack J, Ross T, Bowman JP (2013) Characterisation of the transcriptomes of genetically diverse *Listeria monocytogenes* exposed to hyperosmotic and low temperature conditions reveal global stress-adaptation mechanisms. PLoS One 8(9):e73603. 10.1371/journal.pone.0073603

Elena SF, Lenski RE (2003) Evolution experiments with microorganisms: the dynamics and genetic bases of adaptation. Nat Rev Genet 4(6):457–469. 10.1038/nrg1088

Erickson Keesha E, Otoupal Peter B, Chatterjee A (2017) Transcriptome-Level Signatures in Gene Expression and Gene Expression Variability during Bacterial Adaptive Evolution. Msphere 2(1):e00009–00017. 10.1128/msphere.00009-17

Fang C, Shan Y, Cao T, Xia Y, Xin Y, Cheng C, Song H, Li X, Fang W (2016) Prevalence and virulence characterization of *Listeria monocytogenes* in chilled pork in Zhejiang Province, China. Foodborne Pathog Dis 13(1):8–12. 10.1089/fpd.2015.2023

Gandhi M, Chikindas ML (2007) Listeria: A foodborne pathogen that knows how to survive. Int J Food Microbiol 113 (1):1–15. 10.1016/j.ijfoodmicro.2006.07.008

Gray Michael J, Zadoks Ruth N, Fortes Esther D, Dogan B, Cai S, Chen Y, Scott Virginia N, Gombas David E, Boor Kathryn J, Wiedmann M (2004) *Listeria monocytogenes* isolates from foods and humans form distinct but overlapping populations. Appl Environ Microbiol 70(10):5833–5841. 10.1128/AEM.70.10.5833-5841.2004

Harter E, Wagner EM, Zaiser A, Halecker S, Wagner M, Rychli K (2017) Stress survival islet 2, predominantly present in *Listeria monocytogenes* strains of sequence type 121, is involved in the alkaline and oxidative stress responses. Appl Environ Microbiol 83(16):e00827–00817. 10.1128/AEM.00827-17

Hilliard A, Leong D, O’Callaghan A, Culligan EP, Morgan CA, DeLappe N, Hill C, Jordan K, Cormican M, Gahan CGM (2018) Genomic characterization of *Listeria monocytogenes* isolates associated with clinical listeriosis and the food production environment in Ireland. Genes 9(3):171. 10.3390/genes9030171

Hingston P, Chen J, Dhillon BK, Laing C, Bertelli C, Gannon V, Tasara T, Allen K, Brinkman FS, Truelstrup Hansen L (2017) Genotypes associated with *Listeria monocytogenes* isolates displaying impaired or enhanced tolerances to cold, salt, acid, or desiccation stress. Front Microbiol 8(3):369. 10.3389/fmicb.2017.00369

Holm S (1979) A simple sequentially rejective multiple test procedure. Scand J Stat (6):65-70. 10.2307/4615733

Holt K E, Wertheim H, Zadoks R N, Baker S, Whitehouse CA, Dance D, Jenney A, Connor TR, Hsu LY, Severin J, Brisse S, Cao H, Wilksch J, Gorrie C, Schultz MB, Edwards DJ, Nguyen KV, Nguyen TV, Dao TT, Mensink M, Minh VL, Nhu NTK, Schultsz C, Kuntaman K, Newton PN, Moore CE, Strugnell RA, Thomson NR (2015) Genomic analysis of diversity, population structure, virulence, and antimicrobial resistance in *Klebsiella pneumoniae*, an urgent threat to public health. Proc Natl Acad Sci USA 112(27):E3574–E3581. 10.1073/pnas.1501049112

Hu M, Dong Q, Liu Y, Sun T, Gu M, Zhu H, Xia X, Li Z, Wang X, Ma Y, Yang S, Qin X (2023) A meta-analysis and systematic review of *Listeria monocytogenes* response to sanitizer treatments. Foods 12(1):154. 10.3390/foods12010154

Ji S, Song Z, Luo L, Wang Y, Li L, Mao P, Ye C, Wang Y (2023) Whole-genome sequencing reveals genomic characterization of *Listeria monocytogenes* from food in China. Front Microbiol 13(1):1049843. 10.3389/fmicb.2022.1049843

Kang J, Burall L, Mammel MK, Datta AR (2019) Global transcriptomic response of *Listeria monocytogenes* during growth on cantaloupe slices. Food Microbiol 77(9):192–201. 10.1016/j.fm.2018.09.012

Keeney K, Trmcic A, Zhu Z, Delaquis P, Wang S (2018) Stress survival islet 1 contributes to serotype-specific differences in biofilm formation in *Listeria monocytogenes*. Lett Appl Microbiol 67(6):530–536. 10.1111/lam.13072

Kovacevic J, Ziegler J, Wałecka-Zacharska E, Reimer A, Kitts DD, Gilmour MW (2016) Tolerance of *Listeria monocytogenes* to quaternary ammonium sanitizers is mediated by a novel efflux pump encoded by emrE. Appl Environ Microbiol 82(3):939–953. 10.1128/AEM.03741-15

Li W, Guo Y, Cui Q, Ma X, Zhang X, Chui H, Yang X, Chen W, Li X, Xu B, Chen M, Yu B, Chen W, Fu P, Han H, Liu J, Zhang L (2023) Whole-Genome Sequencing-based characterization of clinical *Listeria monocytogenes* isolates in China, 2013–2019. Foodborne Pathog Dis 20(4):158–168. 10.1089/fpd.2022.0040

Liu D, Lawrence Mark L, Gorski L, Mandrell Robert E, Ainsworth A J, Austin Frank W (2006) *Listeria monocytogenes* serotype 4b strains belonging to lineages I and III possess distinct molecular features. J Clin Microbiol 44(1):214–217. 10.1128/jcm.44.1.214-217.2006

Liu X, Chen W, Fang Z, Yu Y, Bi J, Wang J, Dong Q, Zhang H (2022) Persistence of *Listeria monocytogenes* ST5 in ready-to-eat food processing environment. Foods 11(17):2561. 10.3390/foods11172561

Liu Y, Dong Q, Wang X, Liu B, Yuan S (2021) Analysis and probabilistic simulation of *Listeria monocytogenes* inactivation in cooked beef during unsteady heating. Int J Food Sci Technol 56(5):2282–2290. 10.1111/ijfs.14849

Liu Y, Orsi RH, Gaballa A, Wiedmann M, Boor KJ, Guariglia-Oropeza V (2019) Systematic review of the *Listeria monocytogenes* σ^B^ regulon supports a role in stress response, virulence and metabolism. Future Microbiol 14(9):801–828. 10.2217/fmb-2019-0072

Liu Y, Sun W, Sun T, Gorris LGM, Wang X, Liu B, Dong Q (2020) The prevalence of *Listeria monocytogenes* in meat products in China: A systematic literature review and novel meta-analysis approach. Int J Food Microbiol 312(1):108358. 10.1016/j.ijfoodmicro.2019.108358

Martínez-Suárez JV, Ortiz S, López-Alonso V (2016) Potential Impact of the Resistance to quaternary ammonium disinfectants on the persistence of *Listeria monocytogenes* in food processing environments. Front Microbiol 7 (3):638. 10.3389/fmicb.2016.00638

Maury MM, Tsai YH, Charlier C, Touchon M, Chenal-Francisque V, Leclercq A, Criscuolo A, Gaultier C, Roussel S, Brisabois A, Disson O, Rocha EPC, Brisse S, Lecuit M (2016) Uncovering *Listeria monocytogenes* hypervirulence by harnessing its biodiversity. Nat Genet 48(3):308–313. 10.1038/ng.3501

Moura A, Criscuolo A, Pouseele H, Maury MM, Leclercq A, Tarr C, Björkman JT, Dallman T, Reimer A, Enouf V, Larsonneur E, Carleton H, Bracq-Dieye H, Katz LS, Jones L, Touchon M, Tourdjman M, Walker M, Stroika S, Cantinelli T, Chenal-Francisque V, Kucerova Z, Rocha EPC, Nadon C, Grant K, Nielsen EM, Pot B, Gerner-Smidt P, Lecuit M, Brisse S (2016) Whole genome-based population biology and epidemiological surveillance of *Listeria monocytogenes*. Nat Microbiol 2(2):16185. 10.1038/nmicrobiol.2016.185.

Nelson KE, Fouts DE, Mongodin EF, Ravel J, DeBoy RT, Kolonay JF, Rasko DA, Angiuoli SV, Gill SR, Paulsen IT, Peterson J, White O, Nelson WC, Nierman W, Beanan MJ, Brinkac LM, Daugherty SC, Dodson RJ, Durkin AS, Madupu R, Haft DH, Selengut J, Van Aken S, Khouri H, Fedorova N, Forberger H, Tran B, Kathariou S, Wonderling LD, Uhlich GA, Bayles DO, Luchansky JB, Fraser CM (2004) Whole genome comparisons of serotype 4b and 1/2a strains of the food borne pathogen *Listeria monocytogenes* reveal new insights into the core genome components of this species. Nucleic Acids Res 32(8):2386–2395. 10.1093/nar/gkh562

Nightingale KK, Schukken YH, Nightingale CR, Fortes ED, Ho AJ, Her Z, Grohn YT, McDonough PL, Wiedmann M (2004) Ecology and transmission of *Listeria monocytogenes* infecting ruminants and in the farm environment. Appl Environ Microbiol 70(8):4458–4467. 10.1128/aem.70.8.4458-4467.2004

Omori Y, Miake K, Nakamura H, Kage-Nakadai E, Nishikawa Y (2017) Influence of lactic acid and post-treatment recovery time on the heat resistance of *Listeria monocytogenes*. Int J Food Microbiol 257:10–18. 10.1016/j.ijfoodmicro.2017.06.008

Orsi RH, Bakker HCd, Wiedmann M (2011) *Listeria monocytogenes* lineages: Genomics, evolution, ecology, and phenotypic characteristics. Int J Med Microbiol 301(2):79–96. 10.1016/j.ijmm.2010.05.002

Pightling AW, Lin M, Pagotto F (2014) Draft genome sequence of *Listeria monocytogenes* strain LI0521 (syn. HPB7171), isolated in 1983 during an outbreak in massachusetts caused by contaminated cheese. Genome Announc 2(4):e00729-e00714. 10.1128/genomea.00729-14

Pohl AM, Pouillot R, Bazaco MC, Wolpert BJ, Healy JM, Bruce BB, Laughlin ME, Hunter JC, Dunn JR, Hurd S, Rowlands JV, Saupe A, Vugia DJ, Van Doren JM (2019) Differences among incidence rates of invasive listeriosis in the U.S. FoodNet population by age, sex, race/ethnicity, and pregnancy status, 2008–2016. Foodborne Pathog Dis 16(4):290–297. 10.1089/fpd.2018.2548

Pöntinen A, Markkula A, Lindström M, Korkeala H (2015) Two-Component-System histidine kinases involved in growth of *Listeria monocytogenes* EGD-e at low temperatures. Appl Environ Microbiol 81 (12), 3994–4004. 10.1128/AEM.00626-15

Raser JM, O’Shea EK (2005) Noise in gene expression: origins, consequences, and control. Science 309(5743):2010–2013. 10.1126/science.1105891

Rosenfeld N, Young JW, Alon U, Swain PS, Elowitz MB (2005) Gene regulation at the single-cell level. Science 307(5717):1962–1965. 10.1126/science.1106914

Ryan S, Begley M, Hill C, Gahan CGM (2010) A five-gene stress survival islet (SSI-1) that contributes to the growth of *Listeria monocytogenes* in suboptimal conditions. J Appl Microbiol 109(3):984–995. 10.1111/j.1365-2672.2010.04726.x

Silk BJ, McCoy MH, Iwamoto M, Griffin PM (2014) Foodborne listeriosis acquired in hospitals. Clin Infect Dis 59(4):532–540. 10.1093/cid/ciu365

Skowron K, Wiktorczyk N, Grudlewska K, Wałecka-Zacharska E, Paluszak Z, Kruszewski S, Gospodarek-Komkowska E (2019) Phenotypic and genotypic evaluation of *Listeria monocytogenes* strains isolated from fish and fish processing plants. Ann Microbiol 69(5):469–482. 10.1007/s13213-018-1432-1

Smith A, Hearn J, Taylor C, Wheelhouse N, Kaczmarek M, Moorhouse E, Singleton I (2019) *Listeria monocytogenes* isolates from ready to eat plant produce are diverse and have virulence potential. Int J Food Microbiol 299(6):23–32. 10.1016/j.ijfoodmicro.2019.03.013

Song Z, Ji S, Wang Y, Luo L, Wang Y, Mao P, Li L, Jiang H, Ye C (2022) The population structure and genetic diversity of *Listeria monocytogenes* ST9 strains based on genomic analysis. Front Microbiol 13(12):982220. 10.3389/fmicb.2022.982220

Suzuki S, Horinouchi T, & Furusawa C (2014) Prediction of antibiotic resistance by gene expression profiles. Nat Commun 5(1):5792. 10.1038/ncomms6792

Tahir R, Rabbani M, Ahmad A, Tipu M, Chaudhary M, Jayarao B (2022) Study on the occurrence of *Listeria monocytogenes* in the soil of Punjab Province and its associated risk factors. J Anim Plant Sci 32(1):45–51. 10.36899/JAPS.2022.1.0400

Wemekamp-Kamphuis HH, Sleator RD, Wouters JA, Hill C, Abee T (2004) Molecular and physiological analysis of the role of osmolyte transporters BetL, Gbu, and OpuC in growth of *Listeria monocytogenes* at low temperatures. Appl Environ Microbiol 70(5):2912–2918. 10.1128/AEM.70.5.2912-2918.2004

World Health Organization [WHO] (2018) Listeriosis. https://www.who.int/news-room/fact-sheets/detail/listeriosis. (Accessed 01 March 2023).

Wiktorczyk-Kapischke N, Skowron K, Grudlewska-Buda K, Wałecka-Zacharska E, Korkus J, Gospodarek-Komkowska E (2021) Adaptive response of *Listeria monocytogenes* to the stress factors in the food processing environment. Front Microbiol 12(8):710085. 10.3389/fmicb.2021.710085

Wiktorczyk-Kapischke N, Skowron K, Wałecka-Zacharska E (2023) Genomic and pathogenicity islands of *Listeria monocytogenes*—overview of selected aspects. Front Mol Biosci 10(2):1161486. 10.3389/fmolb.2023.1161486

